# Controlling *Staphylococcus aureus* by 2-[(Methylamino)methyl]phenol (2-MAMP) in a co-culture moderates the biofilm and virulence of *Pseudomonas aeruginosa*

**DOI:** 10.1101/2021.07.03.451022

**Authors:** Gopalakrishnan Thamil Selvan, Brindha Thirunavukkarasu, Nerellapalli Nandini Pravallika, Sahana Vasudevan, Balamurugan Palaniappan, Adline Princy Solomon

## Abstract

*Staphylococcus aureus* and *Pseudomonas aeruginosa* are the most encountered organisms in a polymicrobial chronic wound infection. Production of multiple virulence factors by this duo delays wound healing process. Notably, *P. aeruginosa* displays enhanced virulence in the presence of *S. aureus* by a peptidoglycan sensing mechanism. Thus, novel therapies are imperative to address polymicrobial infections effectively. Previously, it has been suggested that targeting *S. aureus* might be a possible approach to reduce the severity of *P. aeruginosa* in a polymicrobial infection. In this aspect, we have used 2-[(Methylamino)methyl]phenol (2-MAMP), our previously reported QS inhibitor to target *S. aureus* and phenotypically determine the virulence factors of *P. aeruginosa* under this condition. Analysis of major virulence factors of Pseudomonas viz. biofilm, pyocyanin and pyoveridine showed a significant reduction. The competitive index (CI) and relative increase ratio (RIR) were determined to understand the organisms’ interaction in co-culture. Results indicated competitiveness among the strains and on increasing ratios of *S. aureus* cells, co-existence was noticed. Further, as a sensible approach antibiotic – antivirulence drug combinations were tested on co-culture. Significant improvement in the growth inhibition was observed. Our preliminary results presented here would enable further research to address polymicrobial infection in a novel way.

## 1. Introduction

Chronic wounds persist for a long time and remain in the inflammatory phase for too long because of bacterial infections [1, 2]. They are often characterised by biofilm colonization of multispecies organisms. The most encountered organisms among them are *Staphylococcus aureus* and *Pseudomonas aeruginosa* [3, 4]. *S. aureu*s is a significant Gram-positive opportunistic pathogen that causes several diseases from minor skin infections to life-threatening endocarditis, meningitis and osteomyelitis [5]. Whereas, *P. aeruginos*a is a versatile Gram negative bacterium which expresses abundant virulence factors and is one of the leading nosocomial pathogens [6]. In chronic wounds like diabetic foot ulcers, synergistic interactions exhibited by these organisms’ delays the wound healing mechanism. Several studies have shown that *P. aeruginosa* displays enhanced virulence during coinfection with Gram-positive bacteria [7–10]. Moreover, *S. aureus* co-culture interactions offer protection to *P. aeruginosa* from antibiotics like tobramycin, gentamicin and ciprofloxacin [11]. Thus, the opportunistic pathogens growing as mixed biofilms in chronic wounds urge a search for novel diagnostic and treatment strategies.

A few years back, Korgaonkar et al. (2013) [10] reported the mechanism of enhanced virulence of *P. aeruginosa* in co-cultures. The enhanced pathogenesis occurs due to a peptidoglycan sensing mechanism during co-culture with Gram-positive bacteria. Peptidoglycan which contains N-acetylglucosamine (GlcNAc) is shed from Gram-positive bacteria and Pseudomonas responds to that by enhanced production of pyocyanin toxin. This observation is well explained through a drosophila model of infection and murine chronic wound model [10]. Moreover, with their results from mutant strains, they have speculated that targeting *S. aureus* might be a viable approach for reducing the severity of *P. aeruginosa* in polymicrobial infections. To validate this speculation, use of conventional antibiotics is a practical option but downsides the problem of drug resistance as most of the chronic wound *S. aureus* isolates display multidrug resistance (MDR). Alternatively, controlling the pathogens and their virulence factors in a density dependent manner by QS inhibition mediated approach is promising [12]. Inhibition of QS system modifies the gene expression profiles in response to the cell density changes of target bacteria. Interference with QS systems of one pathogen in a co-culture population could modify the pathogenicity and antibiotic resistance of the other [13, 14]. Thus, we have opted to control the virulence of *S. aureus* as it has been clearly shown that in the presence of *S. aureus,* the virulence factors expression of *P. aeruginosa* increases many folds. In our previous studies, we have shown the efficacy of 2[(Methylamino)methyl]phenol (2-MAMP) to inhibit biofilm and virulence of MDR *S. aureus* by targeting the SarA QS pathway [15, 16]. Hence, in the present study we have targeted *S. aureus* by 2-MAMP in a co-culture and assessed the biofilm and virulence factors of *P. aeruginosa*. Further, as a combinatorial approach conventional antibiotic was incorporated in par with 2-MAMP to test growth inhibition efficacy.

## 2. Materials and Methods

### 2.1 Bacterial strains

Standard strains such as *Staphylococcus aureus* ATCC 25923 and *Pseudomonas aeruginosa* ATCC 27853 were used initially to assess the co-culture growth and biofilm. Two clinical isolates, *S. aureus* P1966 and *Pseudomonas aeruginosa* PUS1 (both isolated from a chronic wound infection patient) received from the JSS Medical College, Mysore, India were used. Concerning, the pronounced polymicrobial infections in healthcare settings, more focus was given to clinical isolates in all the assays. All the bacterial strains used in the study were cultured in Tryptone Soya Broth (TSB) [Soyabean Casein Digest Medium) purchased from HiMedia Laboratories, India. The bacterial cultures were incubated at 37 °C in a bacteriological incubator under static conditions in all the assays. The bacterial strains were checked for their purity by streaking onto TSB agar plates and log phase cells were raised from an overnight culture. Glycerol stocks were made with 50% sterile glycerol and stored at −80 °C in an ultralow freezer (NuAire, USA) till use.

### 2.2 Bacterial growth and biofilm

Overnight culture of the strains was obtained from the glycerol stocks by inoculating 10 μl of culture to 5 ml of TSB and incubation at 37 °C. Log phase culture (at 3 h) of the bacterial strains was obtained from the overnight culture. The grown cultures were diluted to 1:50, 1:100 and 1:200 in sterile 1X PBS. The diluted cultures were further serially diluted in 0.85 % saline and plated on TSB agar to standardize the inoculum size as ~10^6^ cfu ml^−1^. Growth and biofilm assessments were made in the presence and absence of QSI compound, 2MAMP at its effective concentration, 1.25 μM [15]. All the assays were conducted in 96-well microtitre plates. To assay monoculture bacteria, 100 μl of the adjusted culture (~10^6^ cfu ml^−1^ in TSB media) was added directly to the wells. Whereas, co-culture conditions were established by adding *S. aureus* and *P. aeruginosa* in the ratios of 100:0, 0:100, 1:1, 1:10, 1:100, 10:1 and100:1 with appropriate microlitres [17]. Compound treatment wells contained 2MAMP at a concentration of 1.25 μM in TSB along with the bacterial cells. After loading the wells, the microtitre plates were incubated for 8 h at 37°C under static conditions in an incubator. Growth was measured at 595 nm in a microtitre plate reader (BioRad i-Mark, Japan).

The biofilm mass was quantified by crystal violet assay. Briefly, planktonic cultures were discarded and the wells of the microtitre plate were washed with 1X PBS to remove the unattached cells. The plate was then air-dried and the wells stained with 0.2 % crystal violet by keeping for 20 min. After 20 min, the dye was removed and wells were washed with 1X PBS twice to remove the excess stain. The bound dye in the wells was eluted with 33% acetic acid and the absorbance of the solution was read at 595nm in a microtitre plate reader (BioRad i-Mark, Japan). To assess the individual bacterial population during growth, planktonic culture from the wells was used for CFU enumeration. For CFU enumeration of biofilm, the attached cells were scraped from the wells and plated onto TSB agar plates after serial dilution. All the plates were incubated for 24 h and the grown colonies were enumerated. CFU enumeration was expressed as cfu ml^−1^ for planktonic and cfu per well for biofilm.

### 2.3 Competitive index (CI) and relative increase ratio (RIR) of *S. aureus* : *P. aeruginosa*

Competitive index (CI) and relative increase ratio (RIR) were calculated to determine the growth behaviour exhibited by the organisms in co-culture. RIR was calculated using the CFU enumeration data from individual growth culture of *S. aureus / P. aeruginosa* at a given time point divided by the same ratio at 0^th^ hour inoculum. CI was calculated using the same method whereas, the CFU enumeration data at a given time point was taken from co-culture growth. A statistically significant difference in CI and RIR indicates competition between the two organisms [18].

### 2.4 Determination of *P. aeruginosa* virulence factors

#### 2.4.1 Pyocyanin

Pyocyanin quantification assay was carried out in the presence and absence of 2MAMP. Briefly, individual and co-cultures of *S. aureus* to *P. aeruginosa* at different ratios (0:100, 1:1, 1:10, 1:100, 10:1 and100:1) were inoculated in 5 ml of King’s medium B (HiMedia Laboratories, India) and incubated overnight at 37 °C. One ml of the overnight culture was pelleted and the supernatant was vortexed with 0.6 ml of chloroform followed by extraction with 1 ml of 0.2 M HCl. 200 μl of the HCl supernatant layer containing the extracted pyocyanin was measured for absorbance at 520 nm with 0.2 M HCl as blank in a multi-mode plate reader (BioTek, USA). The concentration of pyocyanin was calculated by the formula [19]:

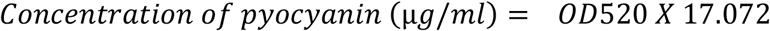

#### 2.4.2 Pyoverdine

Similar to pyocyanin, pyoverdine quantification was carried out in the presence and absence of 2MAMP. Individual and different ratios of *S. aureus* to *P. aeruginosa* co-cultures (0:100, 1:1, 1:10, 1:100, 10:1 and100:1) were incubated in King’s medium B (HiMedia Laboratories, India) for 48 h. The grown cell culture in a microtitre plate was excited at 405 nm and the emitted fluorescence was measured at 467 nm in a multi-mode plate reader (BioTek, USA) [20].

### 2.5 Combinatorial effects of 2MAMP and antibiotics in co-culture

Combinatorial activity of 2MAMP with gentamicin was determined in *S. aureus* to *P. aeruginosa* co-culture ratio of 1:1. Briefly, 100 μl of TSB loaded with 2MAMP (1.25 μM) and gentamicin (32, 16, 8 and 4 μg ml^−1^) were added to the wells of a microtitre plate. Diluted log phase culture containing ~10^6^ cfu ml^−1^ (50 μL each) was inoculated into the wells. Compounds untreated wells were taken as positive control. The plates were incubated for 24 h in an incubator and the optical density was read at 595nm in a microtitre plate reader (BioRad i-Mark, Japan). Growth inhibition percentage was calculated by using the formula:

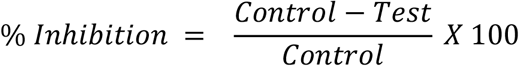

### 2.6 Statistical analysis

All the assays were done in triplicates and statistical analyses were carried out in GraphPad Prism 8.0. The values are expressed as mean ± standard deviation (SD). Differences were considered significant if p≤0.05 with suitable measures mentioned thereof in the results.

## 3. Results

### 3.1 Planktonic growth at mono and co-culture with different bacterial loads

Growth determination was done with individual bacterial population as well as in co-culture. It was observed that the overall growth trend was same at any of the tested co-culture ratios or individual cultures. No significant differences in growth rate were observed in standard and clinical strains as well (Fig. S1). To determine the individual bacterial population, CFU enumeration was done based on the distinct colony morphology between *P. aeruginosa* and *S. aureu*s. The results showed that *P. aeruginosa* outnumbered *S. aureus* in co-cultures compared to monoculture of *P. aeruginosa* (Fig. 1a). However, outcompeting was not seen in *P. aeruginosa* when the proportion of *S. aureus* overpowered *P. aeruginosa* (10:1 and 100:1).

**Fig. 1.**
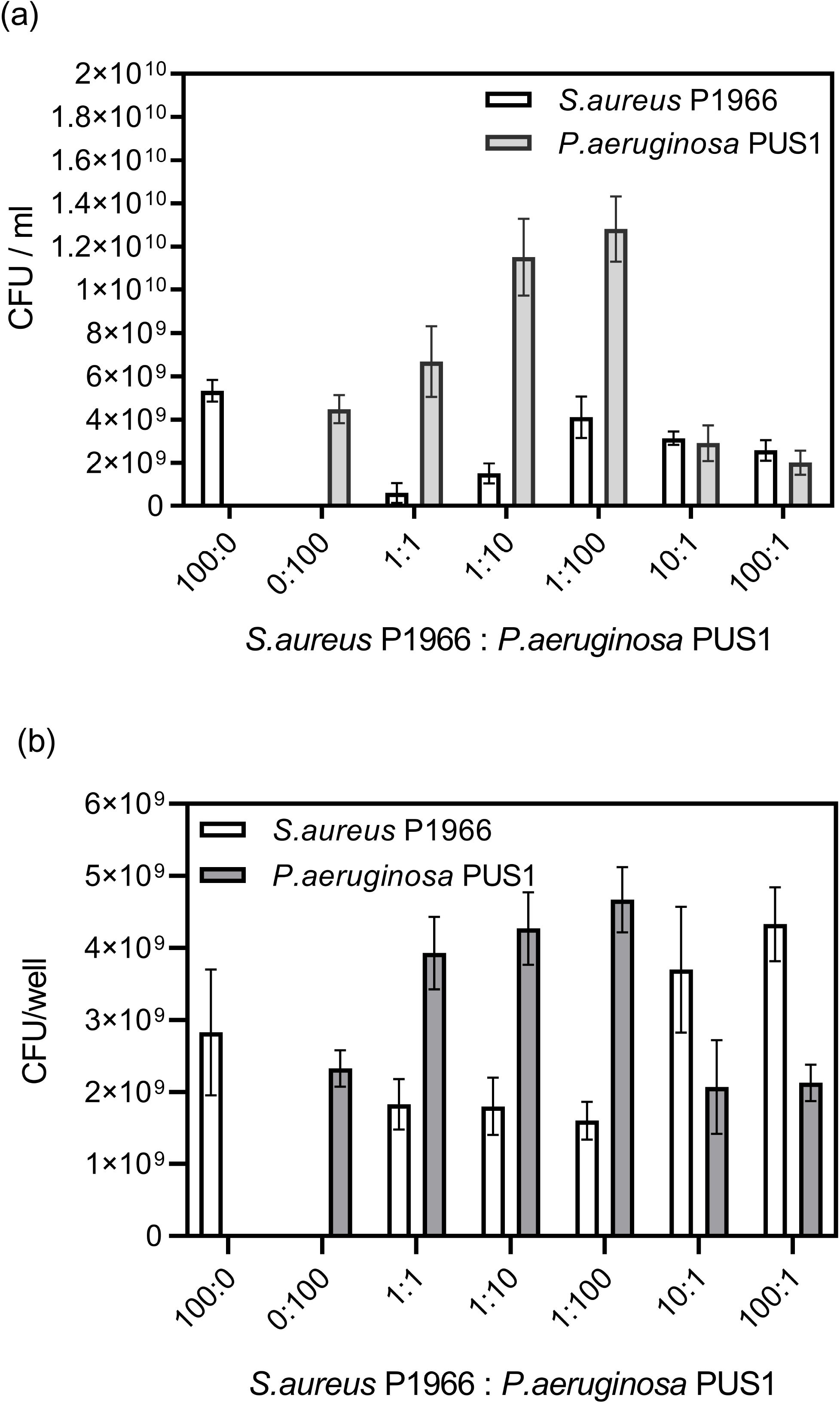
Colony counts from different bacterial loads of *S. aureus* P1966 and *P. aeruginosa* PUS1 at 8 h. a) planktonic bacteria and b) biofilm bacteria. In both planktonic and biofilm cultures, *P. aeruginosa* outnumbered *S. aureus* on equal proportion of cultures (1:1) as well as in increasing *P. aeruginosa* keeping *S. aureus* constant (1:10 and 1:100).

### 3.2 Biofilm formation by mono and co-cultures

Biofilm formation was observed almost in identical levels in the case of standard as well as clinical strains (Fig. S2). No significant difference was seen in the biofilm mass by crystal violet assay in monocultures of *S. aureus* and *P. aeruginosa* and 1:1 ratio of both. At increasing proportions of either *S. aureus* or *P. aeruginosa* (10 and 100), though not significant, a slight variation in the biofilm mass was observed. On CFU enumeration, *P. aeruginosa* was found to be relatively higher than that of *S. aureus* in the scrapped biofilm and contributes to major biofilm formation. Whereas in the higher cell ratios of *S. aureus*, *P. aeruginosa* population is equal to that of found in monoculture (Fig. 1b).

### 3.3 Competitive index (CI) and relative increase ratio (RIR) of *S. aureus* : *P. aeruginosa*

A significant difference in the CI and RIR denotes competition among the co-existing microbes in a polymicrobial culture. Significant difference (P≤0.01; P≤0.05) in the index values were seen on equal proportion of cultures (1:1) as well as in increasing *P. aeruginosa* keeping *S. aureus* constant (1:10 and 1:100) [Fig. 2]. This suggests competition among the strains and no significant difference (P≥0.05) on increasing ratios of *S. aureus* cells suggests co-existence of the microbes.

**Fig. 2.**
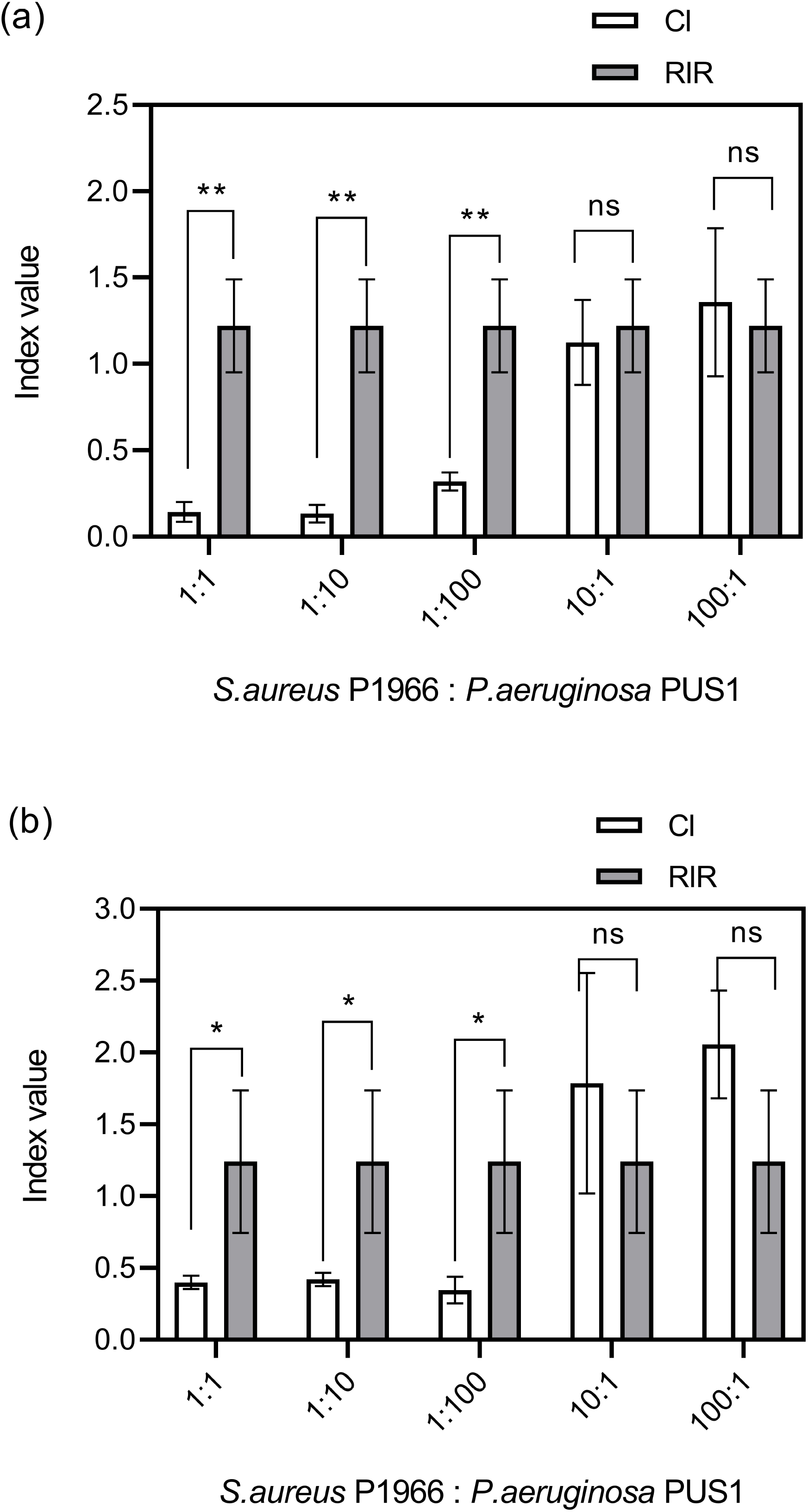
CI and RIR generated from the cfu data of mixed and individual inoculations respectively for *S. aureus : P. aeruginosa*. a) Planktonic growth and b) biofilm growth. Competition between the strains was seen on equal proportion of cultures (1:1) as well as in increasing *P. aeruginosa* keeping *S. aureus* constant (1:10 and 1:100). Statistical significance was determined by an unpaired t-test (**P≤0.01; *P≤0.05; ^ns^P≥0.05).

### 3.4 Effect of 2MAMP against *S. aureus* and *P. aeruginosa* biofilm

Biofilm is considered as one of the virulence factors in an infecting pathogen. Especially in mixed biofilm, one of the major problems is the decreased susceptibility to various classes of antibiotics making it difficult to eradicate. 2-MAMP significantly inhibited the biofilm of *S. aureus* at a concentration of 1.25 μM. It was also confirmed that at the same concentration of 2-MAMP or any other increasing / decreasing drug concentrations *P. aeruginosa* biofilm was not affected (P≥0.05). In case of co-culture ratios, significant inhibition of biofilm (P≤0.0001; P≤0.001) was observed on treatment with 2-MAMP (Fig. 3). On CFU enumeration of the biofilm adhered cells, reduction in cell population of both the organisms were observed on treated samples (Table 1).

**Fig. 3.**
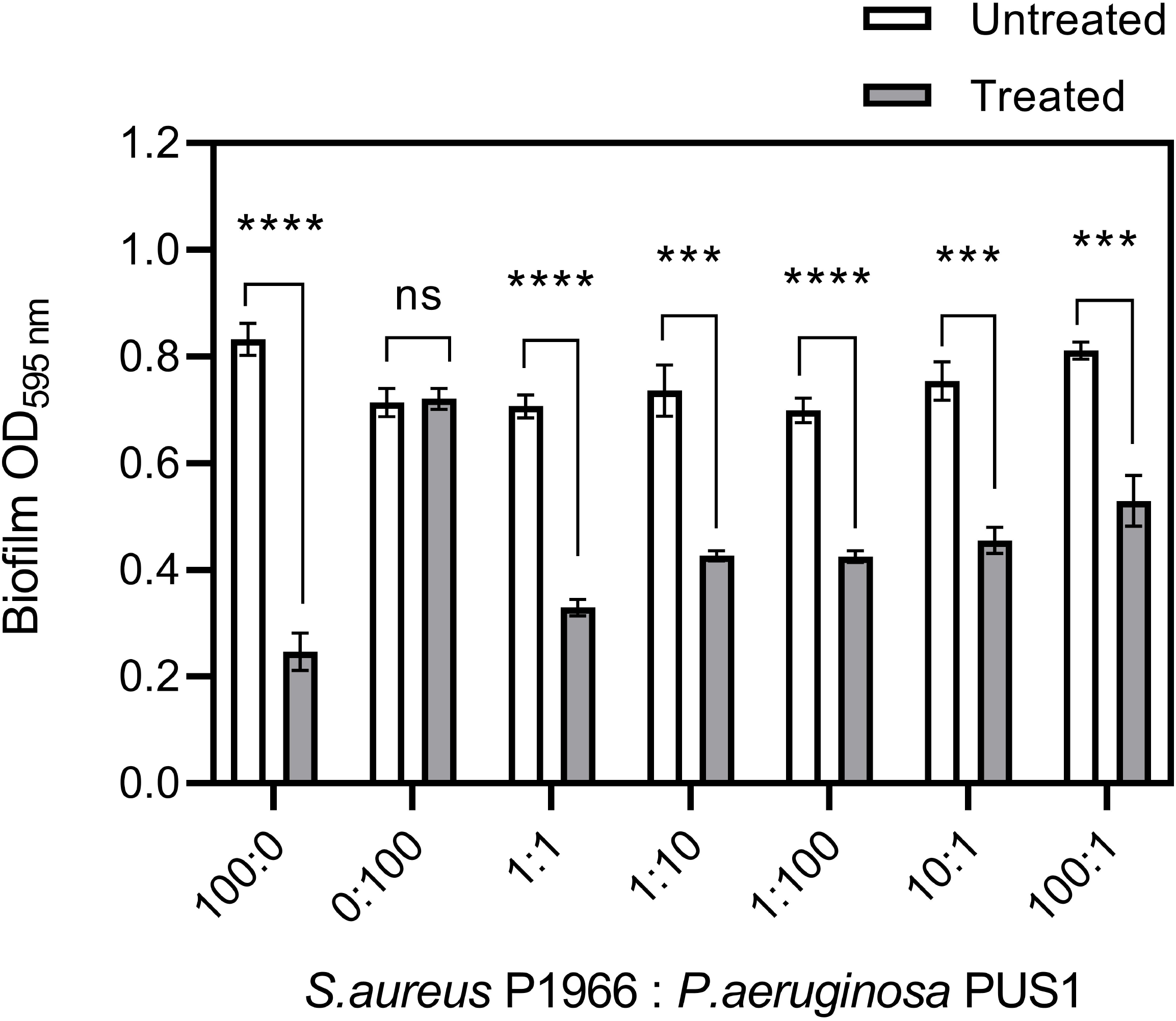
Effect of 2MAMP on biofilm by different bacterial loads of *S. aureus* P1966 and *P. aeruginosa* PUS1. Decreased biofilm mass was noticed in the treated wells. Statistical significance was determined by an unpaired t-test (****P≤0.0001; ***P≤0.001; ^ns^P≥0.05).

### 3.5 Quantification of pyocyanin and pyoverdine on treatment with 2MAMP

Pyocyanin was quantified in the presence and absence of 2-MAMP. As reported earlier, in the present study also we have observed enhanced production of pyocyanin when *P. aeruginosa* is co-cultured with *S. aureus* (Fig. 4a). In the case of 2-MAMP treated samples significant reduction in the levels of pyocyanin was noted compared to the untreated polymicrobial culture. Whatever the ratio may be, reduction is highly significant (P≤0.001; P≤0.01; P≤0.05) and comparable to monoculture of *P. aeruginosa.* The fluorescence exhibited by pyoveridine can be measured at 467 nm. Reduction in the optical density of pyoveridine was noticed in all co-cultures treated with 2-MAMP except for higher proportion of *S. aureus* (100:1) [Fig. 4b].

**Fig. 4.**
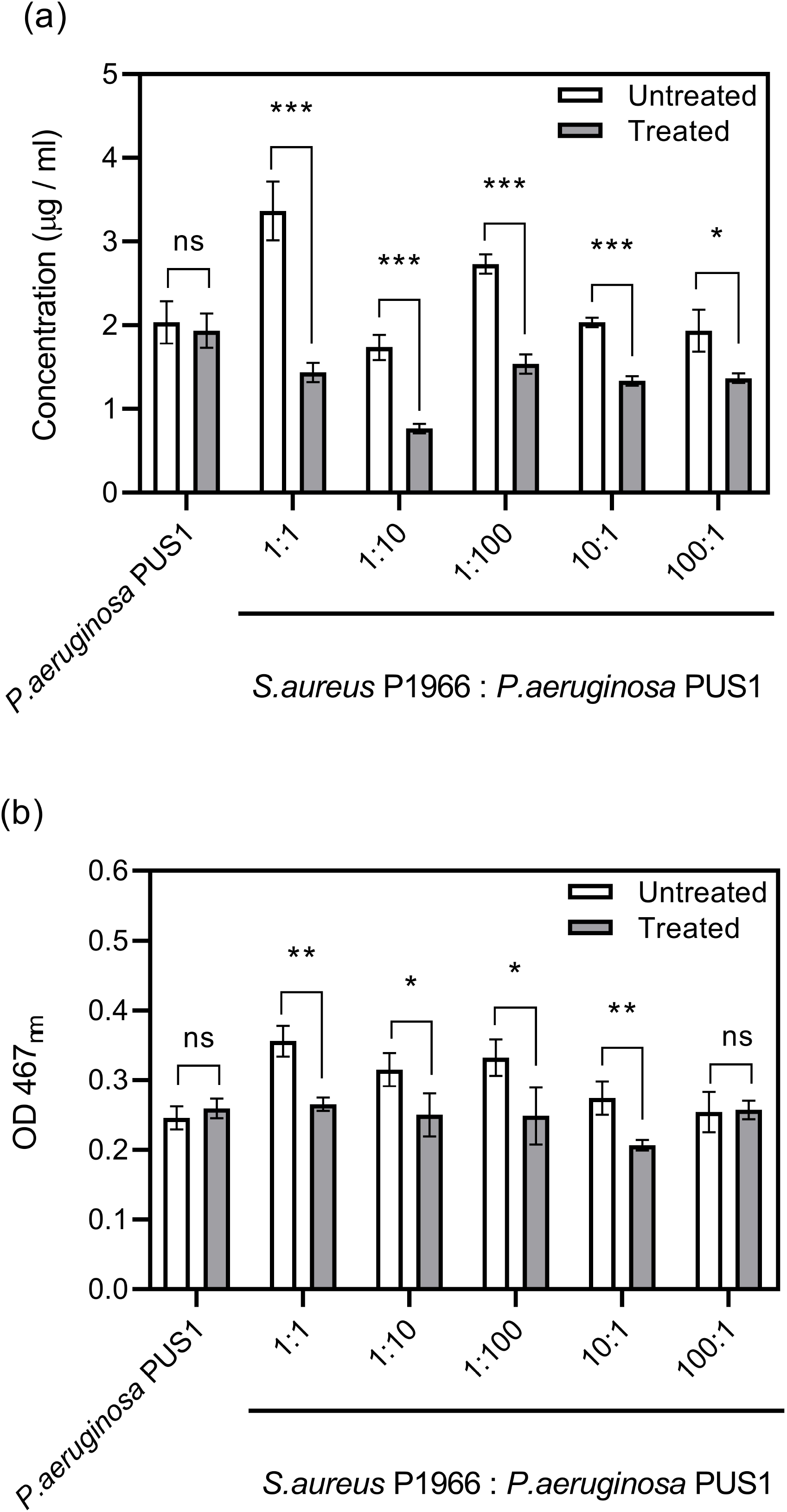
Quantification of a) pyocyanin and b) pyoveridine from untreated and 2MAMP treated co-culture. Reduction in pyocyanin as well as pyoveridine was seen in 2MAMP treated culture. Statistical significance was determined by an unpaired t-test (***P≤0.001; (**P≤0.01; *P≤0.05; ^ns^P≥0.05).

### 3.6 Combinatorial studies

Combinatorial therapy approach is the use of different class of drugs to target drug resistant microbe infections effectively. In the present study, an aminoglycoside antibiotic gentamicin that can act on Pseudomonas as well as Staphylococcus was chosen. At a concentration of 32 and 16 μg ml^−1^ of gentamicin growth inhibition was near to hundred percent (Fig. 5). Whereas, at the sub-inhibitory concentrations of gentamicin alone growth of the co-cultured microbes was inhibited by 50 % and 17 % at 8 and 4 μg ml^−1^ respectively. When gentamicin is combined with 2-MAMP and administered, growth inhibition percentage increased significantly (P≤0.01; P≤0.05).

**Fig. 5.**
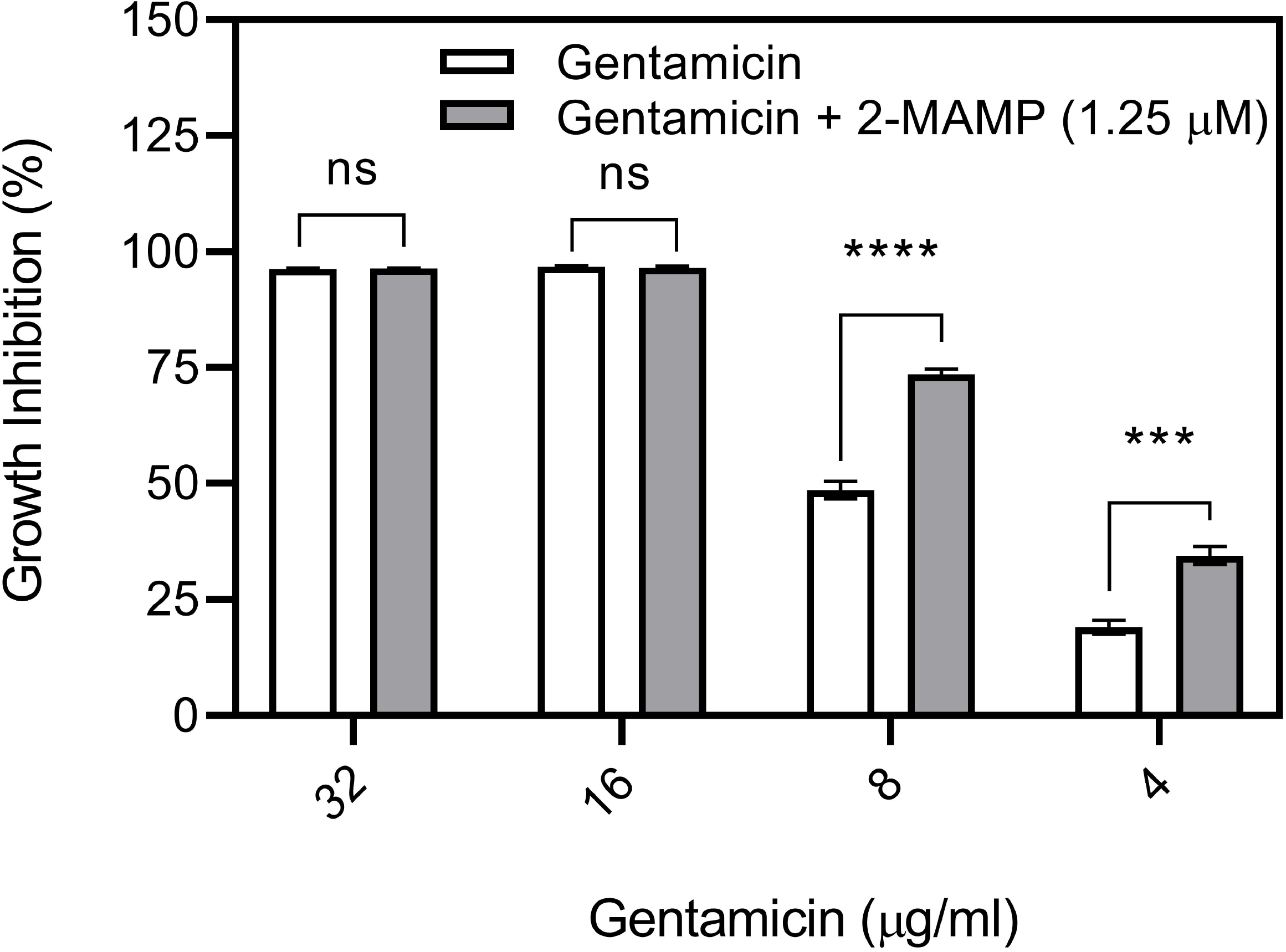
Growth inhibition results of co-cultured *S. aureus* and *P. aeruginosa* by combinatorial approach using gentamicin with 2MAMP. Statistical significance was determined by an unpaired t-test (****P≤0.01; ***P≤0.05; ^ns^P≥0.05).

## 4. Discussion

Microbes associated with polymicrobial infections exhibit competitiveness or synergistic interactions. Such interactions are achieved by intra as well as inter cell-cell communications [3, 10, 17]. Autoinducer small molecules aid these communications in a cell density-dependent manner and the mechanism is termed as quorum sensing (QS) [21]. The QS regulators modulate the enhancement of virulence gene expressions [22]. Besides enhanced virulence, the synergistic interactions among bacterial species favours bacterial persistence at the infection site with increased antibiotic tolerance [23]. It has also been shown that *P. aeruginosa* forms SCVs on co-culturing with *S. aureus* that favours increased survival and antimicrobial tolerance [11].

Novel therapeutic approaches of treating polymicrobial infections are much needed in the present scenario of antibiotic resistance concerns. It is ideal to target a virulence factor that is conserved between different pathogens by using a single anti-virulence drug. However, such targets are not identified between *S. aureus* and *P. aeruginosa* at the present stage. Hence the following strategies would be more practical (i) targeting different strains with different antivirulence drugs (ii) targeting one strain with one antivirulent drug and thereby the virulence of other is modulated and (iii) Combinatorial approach of using antivirulent-antibiotic drugs to control infection. In the present study we primarily focussed on the second approach.

In a polymicrobial infection, one organism influences the growth and biofilm of other. Planktonic growth determinations showed *P. aeruginosa* outnumbering *S. aureus* in co-cultures (Fig. 1a). Similar to planktonic growth, more *P. aeruginosa* was noted compared to *S. aureus* in co-culture in the case of biofilm growth (Fig. 1b). Our results were in corroboration with a study by Baldan et al. 2014 with the isolates from a cystic fibrosis patient [24]. Filkins et al. 2015 showed that *S. aureus* and *P. aeruginosa* coexist initially whereas on extended co-culture, *P. aeruginosa* reduces *S. aureus* viability [25]. Multiple mechanisms have been proposed like siderophore production, genetic, environmental and nutritional factors to explain this behaviour. The most intriguing and principal mechanism is *P. aeruginosa* driven shift of aerobic to fermentative respiration in *S. aureus* [25]. *P. aeruginosa* is capable of lysing *S. aureus* by an antistaphylococcal agent, 2-heptyl-4-hydroxyquinoline N-oxide [26] and use the lysed cells as an iron source [27, 28].

Competitive index (CI) and relative increase ratio (RIR) indicated competition among *P. aeruginosa* and *S. aureus* (Fig. 2). In a polymicrobial infection, competition for the niche can happen through the production of antimicrobials and toxins. For instance, *P. aeruginosa* modifies the composition of microbial community as well as effectuate host killing by this process [10]. In co-culture experiments, *P. aeruginosa* reduced *S. aureus* viability [25]. Sometimes synergy between the coexisting microbes in a polymicrobial infection can happen where the combined effect will be worse than individual microbe effect [21]. Synergism between microbes enhances their survival in the presence of antimicrobials. DeLeon et al. 2014 have observed synergistic effects of *S. aureus* in a co-culture while assessing tolerance to the antibiotics gentamicin and tetracycline [17]. In the present study, on increasing *S. aureus* cells (10:1 and 100:1) co-existence was noted (Fig. 2). A previous study on cystic fibrosis chronic lung infection reported that the adapted strains of *P. aeruginosa* were less capable of outcompeting *S. aureus* when cultured in agar plates. Later, this was found to be because of a commensal-like interaction between the two [11].

As we proposed here, the biofilm and major virulence of *P. aeruginosa* were assessed while targeting *S. aureus* with 2-MAMP. Decreased biofilm mass and cell counts in biofilm (Fig. 3 and Table 1) bestowed positive hope on our tackling approach. Pyocyanin is a redox-active secondary metabolite virulence factor produced by *P. aeruginosa*. The signalling pathways for pyocyanin production and biofilm formation are inter-related [29]. Moreover, enhanced production of the virulence factor pyocyanin during exposure to cell wall fragments of Gram-positive bacteria has been observed [10]. Therefore, quantifying pyocyanin production can give good insights. A significant reduction in pyocyanin and pyoverdine was noted (Fig. 4). Pyocyanin play as a respiratory inhibitor in *S. aureus* forcing them to form small colony variants (SCVs) [30]. Pyoverdine is a fluorescent siderophore produced by *P. aeruginosa* and an important virulence factor required for pathogenesis during infections.

Here we proposed that 2-MAMP would control *S. aureus* by its QS inhibitory activity to maintain a low cell density. This would further limit *P. aeruginosa* numbers as well as reduce the virulence. This speculation is supported by the reduced *P. aeruginosa* numbers observed in 1:10 and 1:100 co-culture ratios (Table 1) where *S. aureus* cell density is maintained extremely low in the initial inoculum. With our preliminary results, we have clearly shown *P. aeruginosa* exhibited reduced adherence and virulence in a co-culture when the partner *S. aureus* is targeted. Furthermore, combinatorial approach with conventional antibiotic enhanced the growth inhibition (Fig. 5). Further, studies on the virulence gene expression levels of *P. aeruginosa,* while targeting *S. aureus* in a co-culture will add more insights on the mechanism. Thereby, through our present study we propose a novel way of addressing polymicrobial cultures which can further be escalated to invertebrate and mammalian infection models towards designing of therapeutic strategies.

## Acknowledgements

We would like to thank: Dr M. N. Sumana (Professor and Head of Department, Department of Microbiology, JSS Medical College, Mysore, India) for providing the clinical isolates; the management of SASTRA Deemed University for providing the infrastructure and lab facilities to complete the research work.

## Author contributions

**GTS, BT, NNP and SV:** Methodology, Validation, Writing - Original Draft. **BP and APS** Conceptualization, Investigation, Supervision, Writing - Review & Editing.

## List of tables

**Table 1.** Cfu enumerated from the scrapped cells of treated and untreated cultures

## Supplementary information

## Supplementary figures

**Fig. S1.** Growth curves of monoculture and co-culture at different bacterial loads. a) *S. aureus* ATCC 25923 to *P. aeruginosa* ATCC 27853 and b) *S. aureus* P1966 to *P. aeruginosa* PUS1. Growth trend was same irrespective of the different ratios of bacterial culture.

**Fig. S2.** Biofilm formation by mono and co-culture at different bacterial loads at 8 h. a) *S. aureus* ATCC 25923 to *P. aeruginosa* ATCC 27853 and b) *S. aureus* P1966 to *P. aeruginosa* PUS1. No significant difference in biomass was observed in the different ratios of bacterial culture.

