## Supplementary figures and images for "Controlling *Staphylococcus aureus* by 2-[(Methylamino)methyl]phenol (2-MAMP) in a co-culture moderates the biofilm and virulence of *Pseudomonas aeruginosa*"

### Figure S1

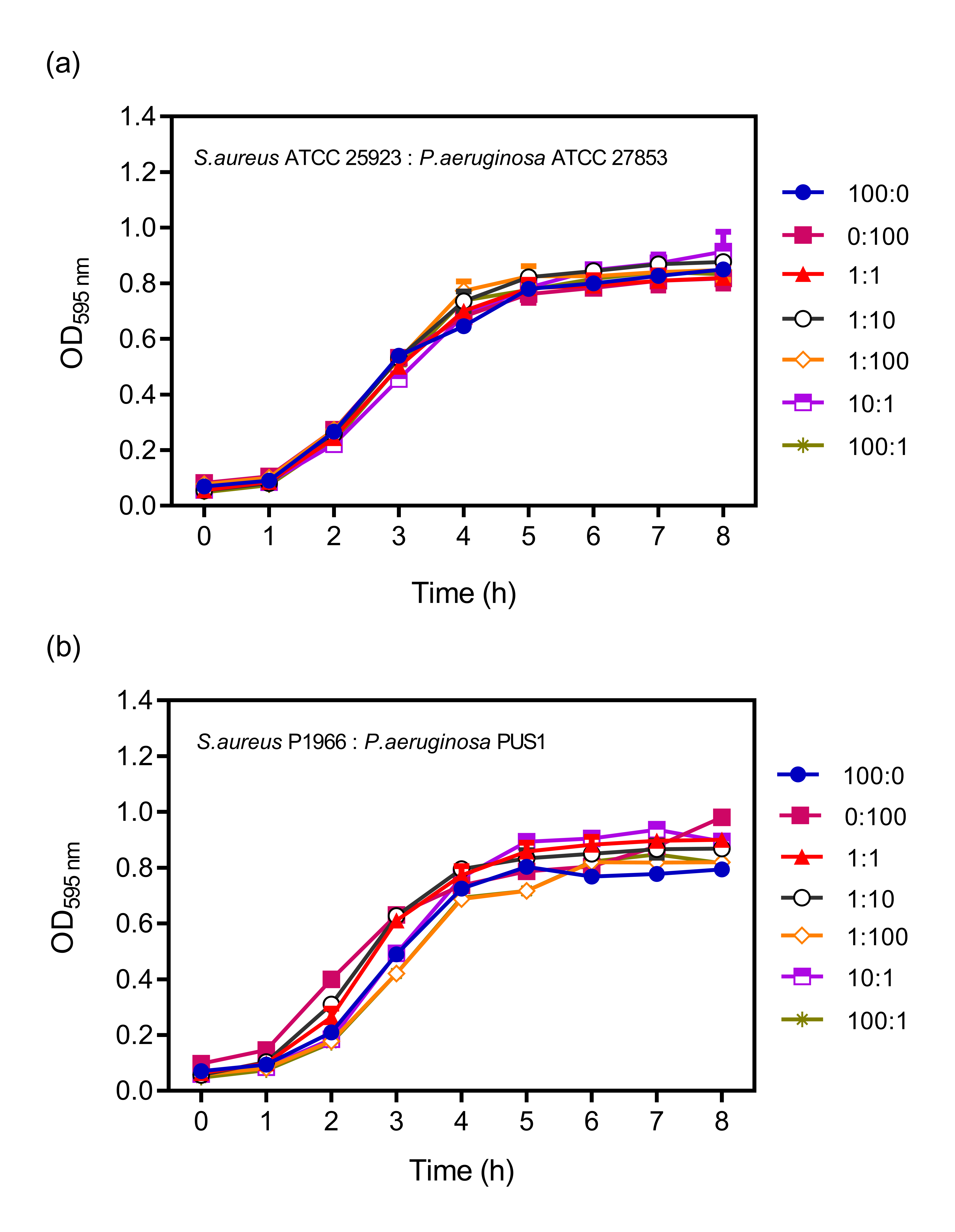

### Figure S2

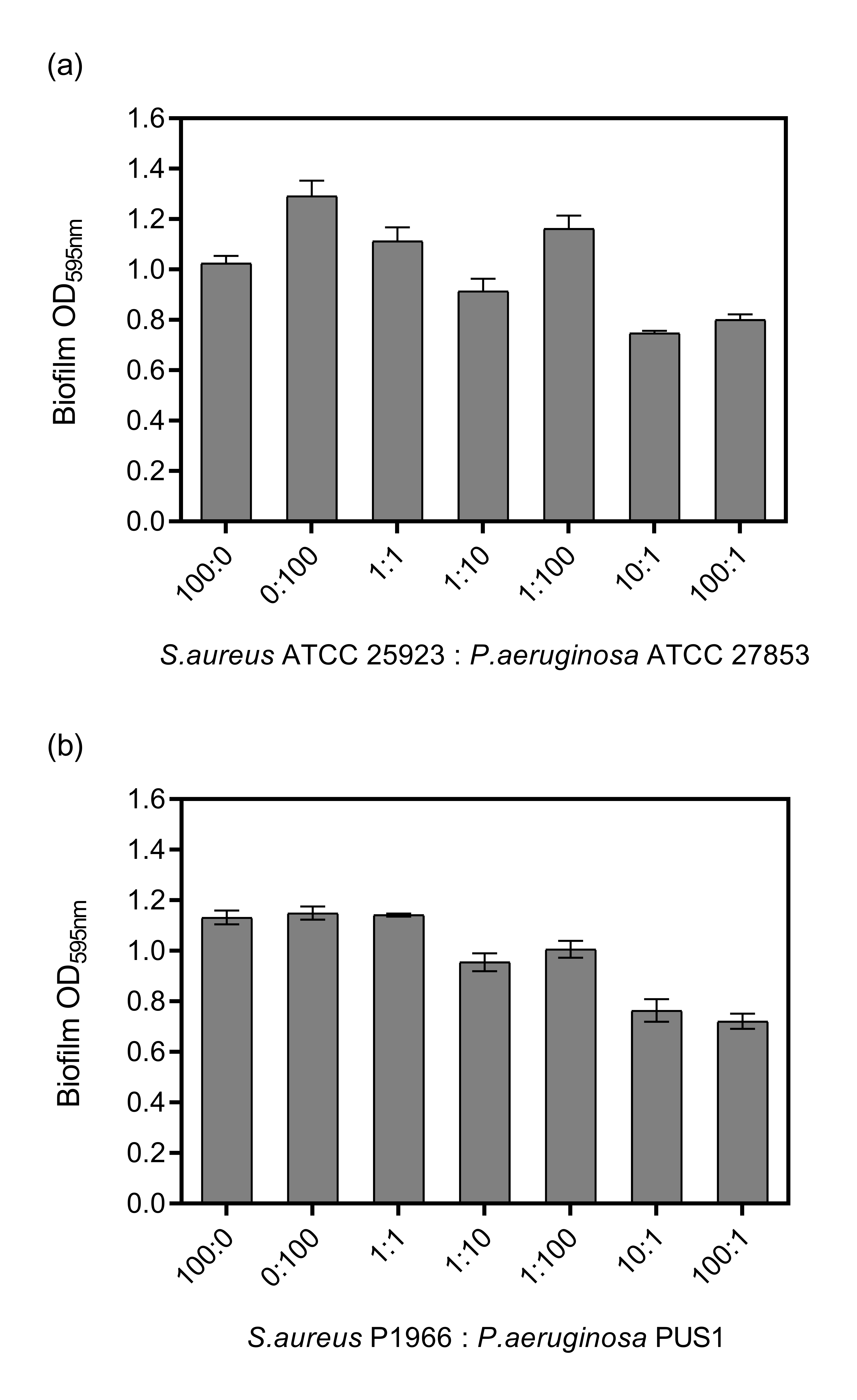
