## Supplementary material for "Controlling *Staphylococcus aureus* by 2-[(Methylamino)methyl]phenol (2-MAMP) in a co-culture moderates the biofilm and virulence of *Pseudomonas aeruginosa*": Table 1

**Table 1.** Cfu enumerated from the scrapped biofilm cells of treated and untreated cultures

| Ratios | Untreated ($x\ 10^9$ cfu/well ± SD) | | 2MAMP Treated ($x\ 10^9$ cfu/well ± SD) | |
|---|---|---|---|---|
| | *S. aureus* P1966 | *P. aeruginosa* PUS1 | *S. aureus* P1966 | *P. aeruginosa* PUS1 |
| 100:0 | 3.1 ± 0.3 | - | 1.8 ± 0.3 | - |
| 0:100 | - | 2.4 ± 0.7 | - | 2.1 ± 0.2 |
| 1:1 | 2.2 ± 0.4 | 4.5 ± 0.5 | 1.5 ± 0.4 | 2.3 ± 0.6 |
| 1:10 | 2.3 ± 0.6 | 4.1 ± 0.8 | 1.4 ± 0.3 | 2.3 ± 0.5 |
| 1:100 | 2.7 ± 0.5 | 4.4 ± 0.6 | 1.5 ± 0.3 | 2.5 ± 0.7 |
| 10:1 | 4.4 ± 0.7 | 2.1 ± 0.4 | 2.8 ± 0.3 | 1.8 ± 0.6 |
| 100:1 | 5.2 ± 0.5 | 2.2 ± 0.7 | 3.7 ± 0.4 | 2.0 ± 0.5 |
